# Competition Between Memory Updating and Differentiation Emerges from Intrinsic Network Dynamics

**DOI:** 10.1101/2025.10.31.685486

**Authors:** Julia Pronoza, Nina Liedtke, Marius Boeltzig, Ricarda I. Schubotz, Sen Cheng

## Abstract

When an event occurs that is similar to a previous experience, the original episodic memory can be modified with new information (updating) or a new memory can be encoded (differentiation). Prediction errors, the deviation between expected and actual stimuli, are believed to mediate the competition between updating and differentiation, but the underlying mechanisms remain unclear. Here, we present a new analysis of experimental studies (Boeltzig, Liedtke, & Schubotz, 2025; Liedtke et al., 2025) that examine recognition memory and cued recall of similar conversations. The original version was recognized more confidently than the modified version, and the recognition confidence for modified versions showed a U-shaped dependence on the prediction error. Furthermore, the larger the prediction error, the more frequently participants retrieved two versions during cued recall. To account for these results, we propose a computational model based on a modified Hopfield network, which encodes the original and modified versions sequentially and weights the encoding of new patterns by the prediction error. The model shows that (1) similar new memories interfere with previous ones (updating) while dissimilar ones are stored separately (differentiation), (2) interference from similar representations leads to reduced memory accuracy and lower-confidence recognition, and (3) the encoding weight must be modulated by the prediction error to account for the experimental data. Our modeling results show that prediction-error-driven competition between updating and differentiation can emerge from intrinsic network dynamics alone.

## Introduction

Imagine driving your usual route to work and noticing a construction sign on the left side of the street. A few days later, you drive the same way, but now see the sign on the right. Was the sign always on the right and you misremember? Or has it actually been moved? These everyday experiences exemplify the dynamic nature of episodic memory: memories are not static snapshots of past experiences, but dynamic, reconstructive representations that can change over time and incorporate new information (Exton-McGuinness et al., 2015; Lee, 2009). When encountering two versions of the same event, the brain can either modify the memory of the first version with information from the second (updating) or retain both separately (differentiation) (Brunec et al., 2020). Prediction errors (PEs), the mismatch between expected and actual outcomes, are thought to play a central role in mediating memory updating (Rescorla, 1972; Sinclair et al., 2021).

Existing studies have examined divergent effects of PEs on memory updating, showing that they can either weaken existing memories (Forcato et al., 2007; Wichert et al., 2013) or promote the integration of new information (Siestrup & Schubotz, 2023; Siestrup et al., 2022; Sinclair & Barense, 2018). However, it remains unclear whether these outcomes reflect distinct processes or variations of a shared updating mechanism. To explore this question, models that capture how PE magnitude influences the interaction between competing memory traces can provide valuable insights.

The predictive coding (PC) framework offers one such account, proposing that the brain continuously adjusts internal models to minimize PEs (Lewis & Bastiaansen, 2015; Millidge et al., 2022; Rao & Ballard, 1999). From this perspective, PEs signal the need to adapt memory, either by modifying existing representations or creating new ones. These processes are further modulated by the precision, or certainty, of the prediction (Feldman & Friston, 2010; Kanai et al., 2015), such that a violation of a very strong prediction elicits a higher PE than a weak prediction does (Henson & Gagnepain, 2010). When an event is largely consistent with a generated prediction, thus evoking a small PE, neural responses are reduced due to increased predictability, known as repetition suppression (Auksztulewicz & Friston, 2016). Conversely, large PEs are thought to allocate attention towards the unexpected information, triggering changes in sensory and higher hierarchical structures to optimize future predictions (Feldman & Friston, 2010; Hohwy, 2012; Itti & Baldi, 2009). Gershman et al. (2017) proposed that decisions about updating versus storing new memories are guided by inferences about latent causes in the environment. If two experiences are inferred to arise from the same cause, the brain updates the memory; if they seem distinct, separate memories are encoded. Within this model, PEs drive both associative learning by updating the connectivity of associated stimuli, when PEs are moderate, and serve as segmentation signals that inform the inference of a new latent cause when discrepancies persist and PEs are sufficiently large. While these models offer powerful insights, they rely on assumptions about the brain explicitly computing statistical properties of stimuli and the minimization of PEs as a primary goal. Yet it is unclear whether such computations are always necessary or biologically plausible. Moreover, most models are grounded in associative learning paradigms, such as Pavlovian conditioning, which may not generalize to the complexities and specific features of naturalistic episodic memory.

Recent neuro-computational studies reveal that PEs can be represented directly within local neuronal circuits, supporting memory adaptation without the need for higher-level inference processes (Makkeh et al., 2025; Mikulasch et al., 2023; Schneider et al., 2025). Studies indicate that PEs can induce changes through local synaptic plasticity, consistent with principles of predictive coding (Whittington & Bogacz, 2017) and associative learning (Den Ouden et al., 2009), suggesting that local, biologically plausible mechanisms may govern complex memory dynamics. Such findings align with classic auto-associative network models of memory, where information is stored and retrieved through distributed attractor states (Hakobyan & Cheng, 2021; Hopfield, 1982).

Here, we propose a computational memory model based on a Hopfield network to examine the dynamics of representations of competing memories. In the regular Hopfield network, memories are stored in a batch using the same encoding weights, so that the order of encoding does not impact the quality of the memory. In our model, the network encodes patterns sequentially and the encoding strength of a pattern depends on its difference from previously stored ones, i.e., the PE or prior accuracy. Larger PEs result in larger synaptic weights. We focus specifically on this aspect of the PE and do not consider the effect of the prior precision, i.e., the certainty of the prediction, in modulating the competition between similar memories. This approach allows for the study of memory representation dynamics using stimuli that differ along one continuous dimension of dissimilarity. Specifically, we study how memory representations change depending on the magnitude of the PE and the strength of the original memory.

This model reproduces findings from recent episodic memory studies (Boeltzig, Liedtke, & Schubotz, 2025; Liedtke et al., 2025), which examined how PE magnitude and original memory strength influence memory outcomes. In these studies, memory accuracy was tested for original conversations and versions of the conversations modified to varying degrees. The model captures three key behavioral patterns that are consistent with previous findings: (1) recognition of original versions is overall higher than that of modified ones, and is less affected by the presence of a competing memory, with highest impairment after medium-level PEs; (2) recognition of modified versions follows a U-shaped function across PE levels, with notably reduced accuracy at intermediate levels of PE; and (3) cued recall likelihood of original conversations remains stable across PE levels, while modified versions are rarely recalled following small changes but are increasingly retrieved as differences become more pronounced. Together, our model offers a step towards bridging the gap between inference-based theories and local circuit accounts by showing that complex memory dynamics, including updating, interference, and differentiation, can emerge from the dynamics of a continuous, PE-modulated mnemonic network.

## Methods

To investigate how the PE mediates whether memories are updated or differentiated when similar experiences are encountered, we performed a new analysis of previously published experimental data (Boeltzig, Liedtke, & Schubotz, 2025; Liedtke et al., 2025), and developed a computational model to account for the experimental results.

### Experimental Studies

In Study A, participants listened to a sequence of 30 naturalistic conversations (Fig. 1A), which were presented as auditory stimuli three or five times. On the subsequent day, participants heard the conversations again, but 24 out of 30 were modified to varying degrees, without prior notice (Fig. 1B, left column). One of four prepared versions was chosen as the modification, which comprised low or high surface or gist manipulation. The conversations were always modified in the middle (target) section of the dialogue, such that the beginning and the end were the same as in the original version (Fig. 1A). The following day, recognition memory was tested, where participants were presented with snippets of the two familiar versions (the original and the modification), new modifications of heard originals, and completely new conversations. Participants were asked whether they remembered the conversation snippets with a confidence rating from 1 (definitely new) to 6 (definitely old). Subsequently, cued recall was tested, where participants were cued with the common, unmodified beginning of the conversation and were asked to verbally complete the conversation. In case they remembered more than one version, they were allowed to give more than one response. After the memory test, subjects gave individual difference ratings for the conversations used in this study, ranking the perceived difference between the two presented versions on a scale of 1 (very similar) to 7 (very different) (Fig. 8A). For more details on this study see Boeltzig, Liedtke, and Schubotz (2025).

**Figure 1:**
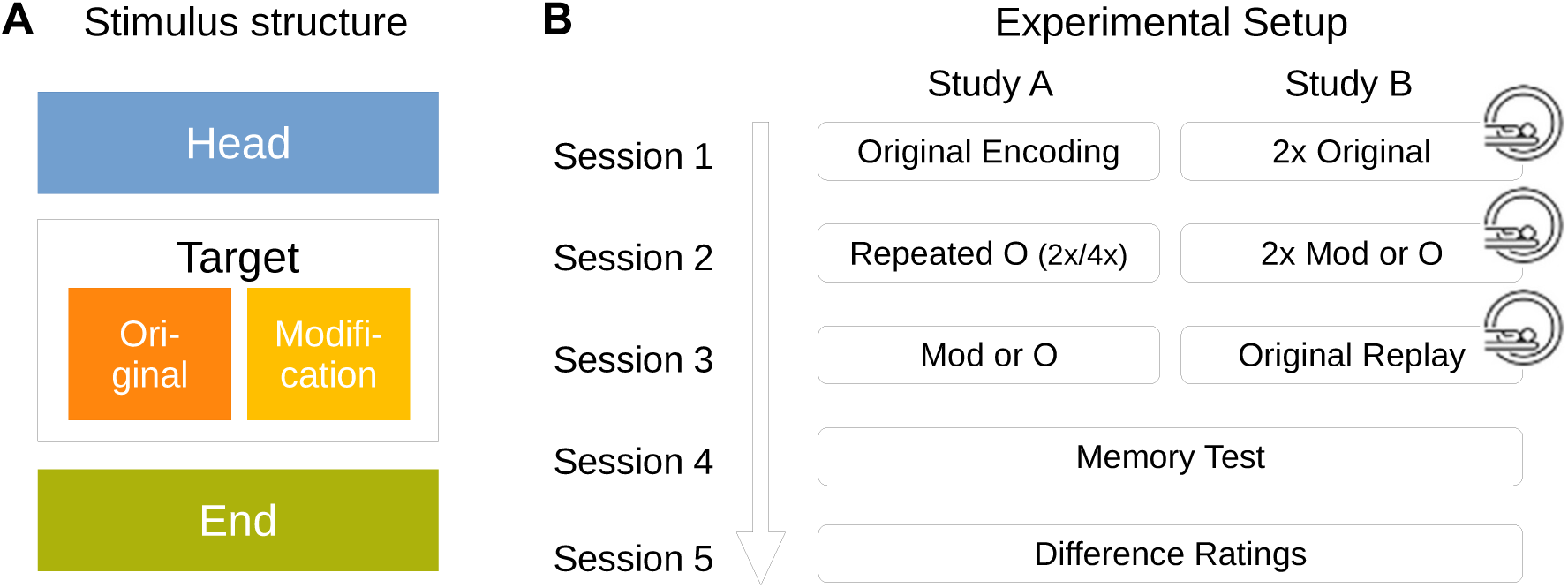
Experimental Design. **A:** The conversations used in both studies consisted of three parts; the common beginning (head), the middle (target), which was presented in the original or a modified version, and a common ending. **B:** In both studies, participants first encoded original followed by modified versions of the conversational stimuli, then carried out a recognition and cued recall test, and finally gave difference ratings for the stimuli. The two studies differed primarily in the frequency of encoding of the two versions. Additionally, sessions 1–3 of study B were conducted in an fMRI scanner, but the data are not analyzed here. Figure adapted from Boeltzig, Liedtke, and Schubotz (2025) and Liedtke et al. (2025).

Study B used the same stimulus material and memory tests as study A, but the study design was altered to enable pre-post comparisons of BOLD signals acquired during stimulus presentations (Fig. 1B, right column). The neuro-imaging data is not analyzed here. Participants listened to each original conversation twice on the first day, the modifications were presented twice in a second session, and then the originals were played once more in a third session. For more details see Liedtke et al. (2025).

In the studies mentioned above, the analyses differentiated between memory of surface and gist modifications. Here, we present the results using only the difference rating obtained from the participants as a measure of the PE magnitude. Furthermore, we also analyze the recall data collected in the aforementioned studies.

#### Cued Recall Analysis

To evaluate the responses given by the participants during cued recall, we used a large language model for sentence embeddings (Reimers & Gurevych, 2020). These embeddings allowed us to quantify the semantic similarity between the responses given by the participants and the presented versions of the conversations using the cosine similarity measure.

### Modeling Framework

Our model of the competition between updating and differentiation of memories is based on the Hopfield network (Hopfield, 1982). The general idea is that this simple model can account for updating and differentiation based on network dynamics as follows: The representations of dissimilar memories form well-separated attractors in the Hopfield network, which can be retrieved individually (differentiation, Fig. 2A). By contrast, two attractors representing very similar memories overlap, fusing into a new, merged representation (updating, Fig. 2B), which makes it impossible to retrieve the memories separately.

**Figure 2:**
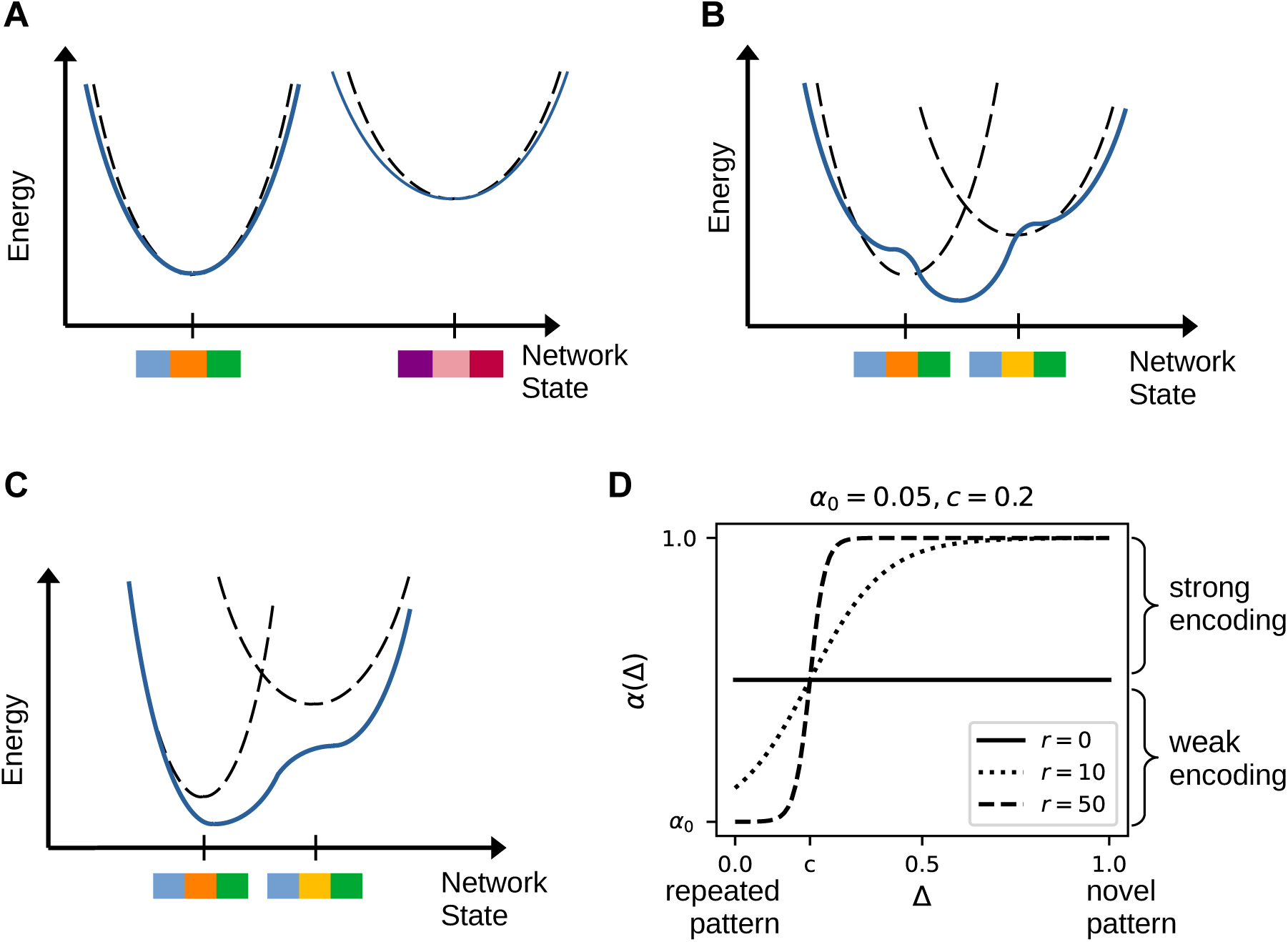
Illustration of interference between attractors in the Hopfield network. Dashed, black lines represent the network energy of individually encoded memories, solid blue lines represent the network energy when multiple memories are stored simultaneously. **A:** The attractors representing two dissimilar patterns (indicated by the barcodes along the horizontal axis) are well separated, leading to differentiation. **B:** Attractors representing similar patterns encoded with similar strength interfere with each other, leading to a merged attractor, i.e., updating of the first pattern. **C:** If one pattern is encoded more strongly, a new attractor results that is biased toward the stronger pattern. **D:** The logistic function used for the encoding strength *α*(Δ). A constant encoding strength is obtained for *r* = 0 (solid line). The larger the parameter r, the more sudden the transition between weak and strong encoding becomes.

The memories of conversations were represented as vectors *p* with 3000 binary entries, these patterns were sequentially encoded in the network weights. Upon presentation of a partial cue, a pattern was retrieved from the Hopfield network through iterative network updates. A recognition memory decision was made based on the difference between the cue pattern and the retrieved patterns (Hakobyan & Cheng, 2021). We studied recognition memory and cued memory recall as a function of the difference between original and modified patterns, as well as the encoding strength of the original pattern.

### Encoding

The standard Hopfield network encodes all patterns the same way, regardless of the order in which the patterns were presented and the number of presentations. To allow these aspects to influence the encoding of patterns, we altered memory encoding in the Hopfield network by making it dependent on the similarity of a new pattern *p^µ^* to previously stored patterns as follows. First, using *p^µ^* as a cue, a pattern 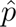 was retrieved by the network (Fig. 3A), which can be thought of as a prediction of the model. Therefore, the distance between the two patterns

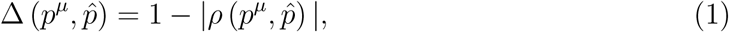

**Figure 3:**
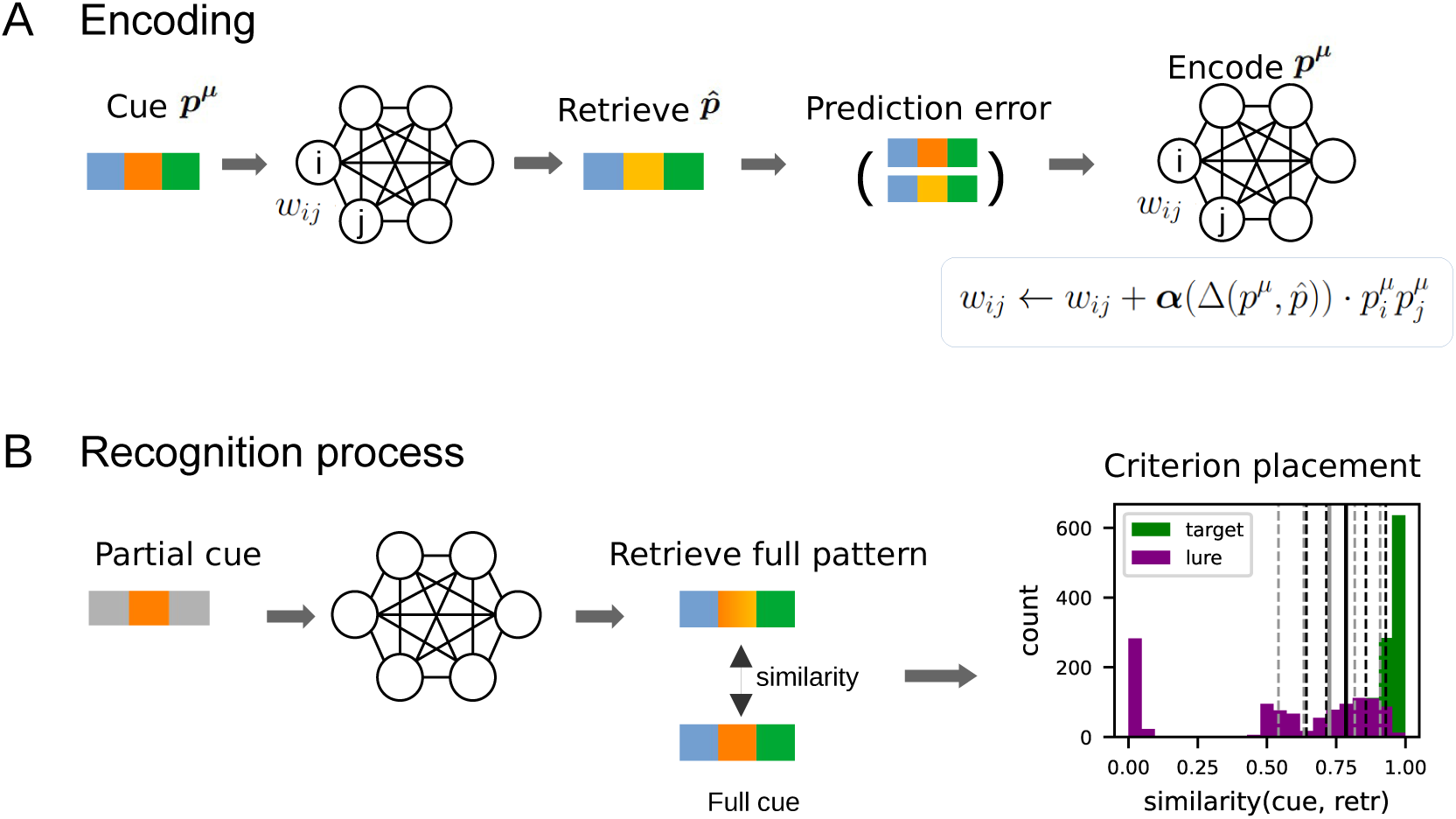
Modeling framework consists of PE–modulated encoding and confidence-based recognition. **A**: During encoding, target patterns to be stored are presented to the network as cue. Upon convergence, the retrieved pattern is compared to the cue to make a prediction, and the distance between the two patterns then represents the PE Δ. The new pattern is stored in the network with an encoding strength *α*(Δ). **B**: During the recognition test, the studied patterns are partially presented to the network, which then performs a cued recall. The retrieved pattern is compared to the cue pattern to make a recognition judgment. A confidence rating is given on a scale of one (low similarity / definitely new) to 6 (high similarity / definitely old) based on evenly placed response criteria. The criterion placement varies between stricter (black lines) and more liberal (gray lines) distributions in different trials. The solid lines indicate the decision threshold *θ*_3_ between old and new.

where *ρ* is the correlation coefficient can be interpreted as a PE. For new patterns that are distinct to previous ones Δ = 1; for exact repetition of a previous pattern Δ = 0. Intermediate values occur, if the current pattern is similar to a stored pattern. Second, the current pattern was stored in the network by updating the network weights with a PE-dependent encoding strength

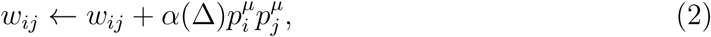

where 0 ≤ *α* ≤ 1. We found that the dependence of the encoding strength on the PE has a profound impact on memory performance. In particular, it determines how interfering patterns shape the resulting attractor state (Fig. 2C). In our simulations, we chose a logistic function (Fig. 2D):

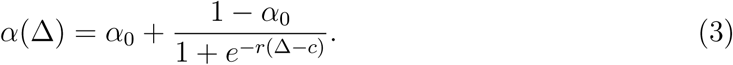

Hence, *α* approaches *α*_0_ for low PEs, which determines how strongly an exactly repeated pattern (Δ = 0) is encoded. The lower *α*_0_, the weaker the encoding of a repeated pattern. With increasing Δ, the encoding strength approaches 1. The larger the midpoint of the curve *c* or the smaller the slope *r*, the larger the PE at which the network encoded the modification differently from the original (Fig. 2D).

### Study Patterns

We stored 52 binary, uncorrelated patterns of length 3000 in the network. Of these, 20 served as baseline memories, providing comparison patterns for the original patterns and partially filling network capacity to make interference effects more evident. Of the remaining patterns, 16 are additionally encoded twice (weak encoding) and 16 four more times (strong encoding). 28 of the 32 repeatedly encoded original patterns were modified to varying degrees and the modifications were encoded in the network subsequently to the original ones. The modified patterns were generated from the originals by flipping the state of a fraction of units (ranging from 0.05 to 0.75) randomly chosen from the middle section of the pattern, corresponding to the target region manipulated in the behavioral study. Increasing the number of flipped units increases the difference between the original and modified patterns. To account for individual levels of memory accuracy in subjects, encoding noise was introduced to all stored patterns by flipping the values of a small fraction of randomly chosen units. The fraction was drawn from a half-normal distribution with mean 0 and standard deviation *σ_enc_*.

### Recognition Memory Test

For the recognition test, retrieval was cued with the middle, i.e., the potentially modified, part of each studied pattern. After the network activity converged, we computed the retrieval similarity, i.e., the correlation *s* = *ρ* (*p,* 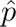) between the retrieved pattern 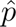 and the full cue *p*. This retrieval similarity was then converted to confidence levels *ζ*, measured on a six-point scale as used in recognition memory experiments. A high retrieval similarity indicates that the cue probably corresponds to a stored pattern, i.e., the cue is an old stimulus (4-6). By contrast, a low retrieval similarity indicates that the cue has no close association with any of the stored patterns and leads to low-confidence responses, i.e., the cue is new (1-3). The numeric conversion is based on the model of Hakobyan and Cheng (2021). We defined *n* = 5 threshold values (Fig. 3B):

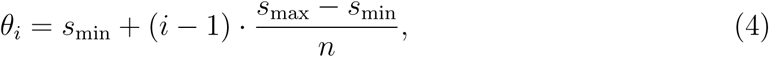

where *i* = 1*,…, n*. The upper boundary was set at *s_max_* = 1 and the lower boundary *s_min_*varied to adjust for individual response characteristics. We successively checked if *s < θ_i_* for *i* = 1*,…,* 5. The first *i* for which this condition held became the confidence level, i.e., *ζ* = *i*, else *ζ* = 6.

Studies have shown that the inclination to give a high-confidence response can vary from person to person (Aminoff et al., 2012; Kantner & Lindsay, 2012), leading to a variance in performance, even if the memory quality is the same. In our model, the larger *s*_min_, the stricter the responses, such that high-confidence old responses become less likely (Fig. 3B). The ideal case is when the confidence thresholds are distributed such that all targets (previously encoded items) are correctly labeled as “old” (confidence level ≥ 3) with a minimum false alarm rate (accept non-encoded items, known as lures, as “old”). We therefore chose a reference *s*_min_ so that the similarities for all target item *s > θ*_3_, which means a confidence *ζ* ≥ 4 (old). To model variations in individual decision-making rigor, we shifted *s*_min_ from trial to trial by a value drawn from a uniform distribution between 0 and 0.1.

## Results

### The Role of the Encoding Strength in Memory Interference and Differentiation

We first studied how the network parameters in Eq. (3) affect memory encoding in the model using the similarity between studied cue and retrieved pattern as an indicator of the encoding accuracy. If *r* = 0 in Eq. (3), patterns are encoded independently of the PE (Fig. 2D, solid line), i.e., original and modification are treated the same and lead to similar encoding outcomes (Fig. 4A). If *r >* 0, the encoding strength is higher for novel stimuli than for similar stimuli (Fig. 2D, dashed lines), resulting in stronger attractors for original patterns than for modified ones. The larger *r*, the larger the difference between the encoding weights of the original and weakly modified versions (Δ *< c*), and the smaller the difference of less similar modifications (Δ *> c*) (Fig. 4A). For a repeated pattern, i.e, Δ(*p^µ^,* 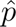) = 0, the encoding strength is *α*_0_. The higher *α*_0_, the more synaptic weight is attributed to the repeated pattern and the more robust this pattern becomes (Fig. 4B, solid lines). Conversely, a competing, weakly encoded memory, as is the case when encoding a modification, becomes less stable, as the strong original attractor absorbs the weak attractor of the modification. Thus, with larger *α*_0_ modifications can only be robustly retrieved after larger PEs (Fig. 4B, dashed lines). A similar effect is observed when increasing the midpoint *c* of the encoding strength function. This parameter moves up the PE value at which the modification is attributed substantially higher encoding strength and can be differentiated from a previously encoded original (Fig. 4C). Another parameter that controls the retrieval accuracy in our model is the encoding noise. It is not related to the encoding strength, but overall deteriorates the encoding of all patterns. Higher values of *σ_enc_* lead to noisier stimuli and results in lower accuracies and higher variances (Fig. 4D).

**Figure 4:**
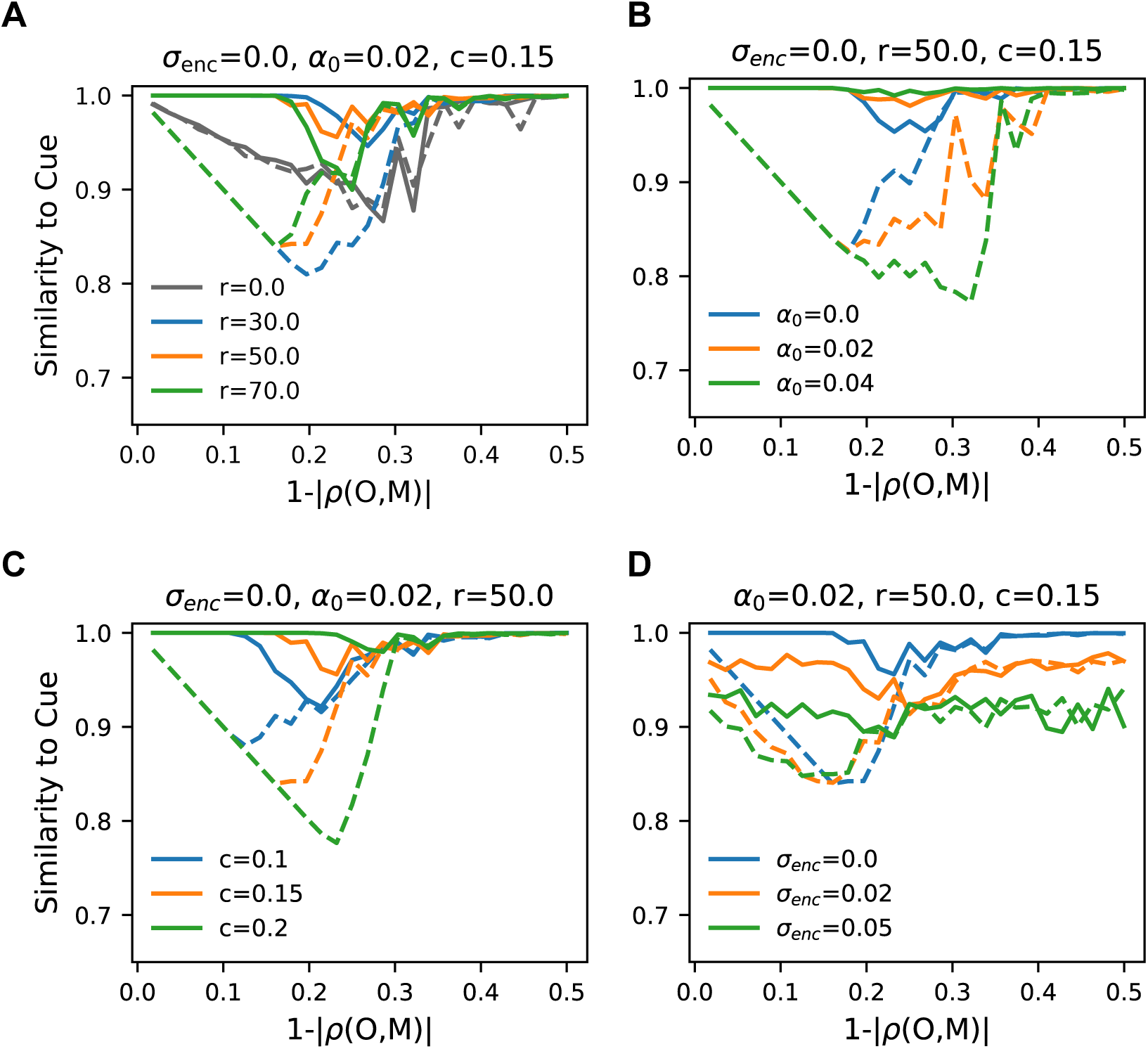
Effects of varying model parameters on retrieval. Solid lines indicate the retrieval similarity to original cue patterns and dashed lines the retrieval similarity to modified cue patterns as a function of the difference between the original and modified versions. **A**: When the slope of the encoding strength function *r* = 0, original and modified patterns are encoded with equal strength *α* and lead to equivalent retrieval curves (gray lines). Positive values of *r* lead to stronger encoding of the original. The larger *r*, the smaller the difference between encoding of the original and large modifications, i.e., 1 − |*ρ*(*O, M*)| *> c*. **B**: The larger the encoding strength of a repeated pattern (*α*_0_), here with three presentations of the original pattern, the more stable the original and the more instable the modification becomes. **C**: Shifting the midpoint *c* of the encoding strength function shifts the range where encoding of the original and modification differ from one another. **D**: Adding noise to the encoded patterns deteriorates encoding of patterns generally.

Next, we asked which features of the model are required to account for the recognition memory results from the experimental studies. We first considered the case of a single encoding of the original and modified versions with constant encoding strength, i.e., *r* = 0 in Eq. (3). In this case, patterns are encoded independently of one another and the parameters *α*_0_ and *c* do not affect the outcome. The similarity between retrieved patterns and cues is a U-shaped function of the distance between original and modified patterns (Fig. 5A), which leads to a U-shape in the confidence rating (Fig. 5C). These results suggest the following interpretation: if original and modification are similar, the network cannot store them as separate attractors and therefore encodes an interference pattern. This pattern is retrieved regardless of whether the network is cued with the original or modified pattern. The more the original and modified patterns differ, the more the interference differs from both, and hence the retrieved pattern is less similar to the cue pattern. As the original and modification become even more dissimilar, this effect reverses and interference is reduced, because both patterns can be stored and retrieved separately. Hence, for similar stimuli the model performs memory updating, i.e., the previously stored memory is altered by the storage of the new memory, while for larger modifications the model performs differentiation, by encoding both patterns as separate memories (Fig. 2).

**Figure 5:**
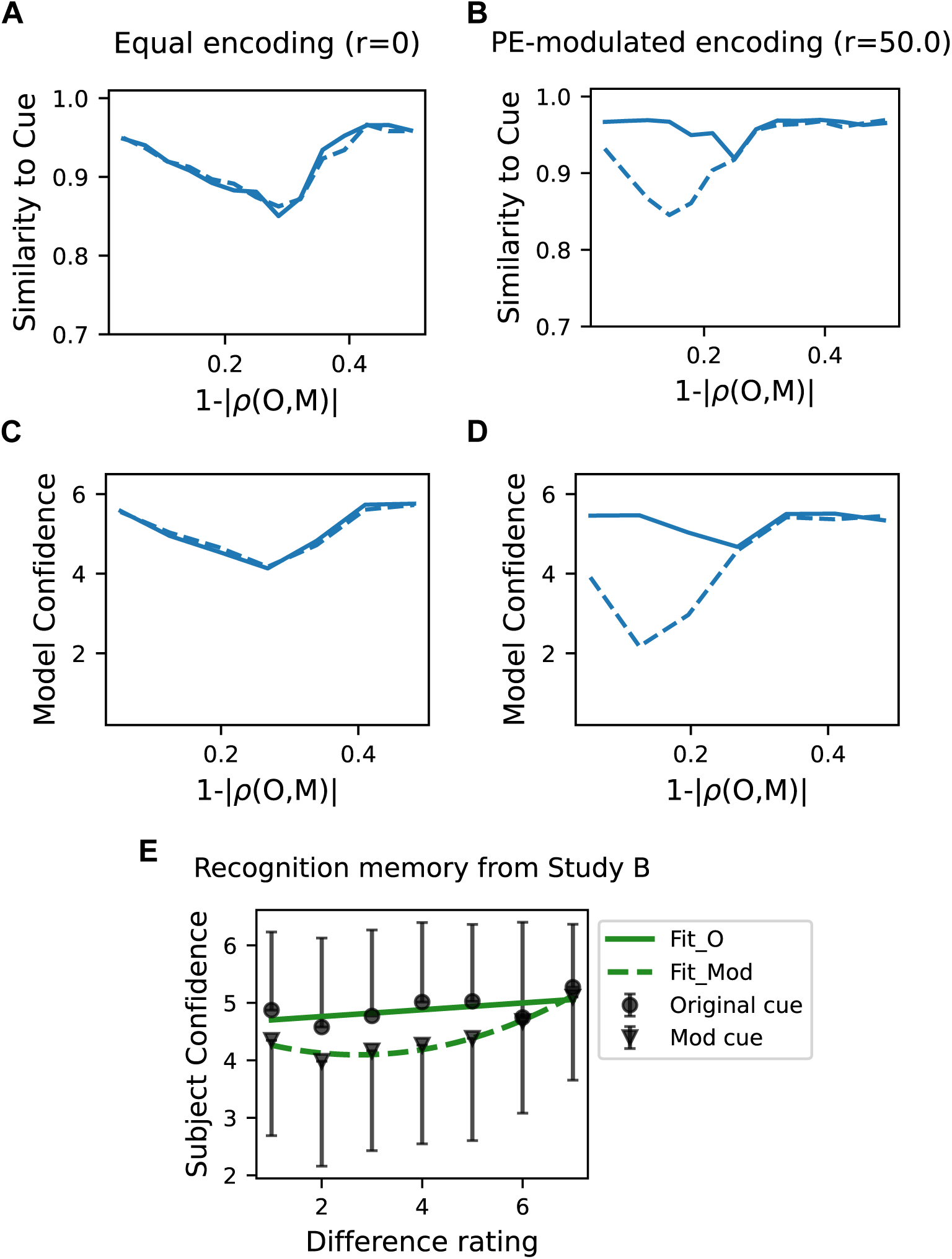
Encoding strength modulated by prediction error is required to reproduce behavioral patterns from Study B. **A**: Encoding patterns with equal encoding strength *α* = const (*r* = 0, Eq. (3)) results in equivalent retrieval curves for original (solid line) and modified (dashed line) versions. The curves are u-shaped due to an interplay between updating and differentiation (see main text). **B**: Modulating the encoding strength with the PE leads to a stronger encoding and robust retrieval of the original and weaker encoding of the modification. For small modifications, retrieval is biased towards the original, resulting in a u-shaped retrieval curve for modified cues. **C**, **D**: Recognition memory confidence rating given by the model for the same simulations as **A** and **B**. By definition (Eq. (4)), the confidence ratings reflect the retrieval similarity of original and modified cues. **E**: Results of the recognition memory test of Study B. Shown are average confidence ratings for recognition of original (dots) and modified (triangles) stimuli as a function of participant rated difference. The U-shaped curve for modified stimuli and higher recognition confidence for original stimuli are explained by the PE-modulated model.

The U-shape of the recognition curve for the modified versions observed in the model is consistent with robust experimental results from both studies (Figs. 5E, 6C). However, in the model recognition for the modification is too good and recognition of the original version should be more robust to the PE. These two shortcomings can be addressed by making the encoding strength dependent on the PE (*r >* 0, Eq. (3)). At this setting, weakly modified patterns, i.e., low PE, are encoded with a small encoding strength *α*, so that the original patterns can still be retrieved faithfully, but cueing the modified version also retrieves the original pattern (Fig. 5B). Therefore, memory confidence remains high for modifications because the retrieved pattern is highly similar to the cue due to the small differences between two versions (Fig. 5D). When the difference between the two versions becomes larger, the encoding strength of the modification increases (Fig. 3 B), leading to a separation of the associated representation. Thereby the resulting interference pattern is biased towards the more strongly encoded original, leading to lower retrieval confidence for the modification compared to the original. Finally, when modifications are sufficiently large, the encoding strength approaches *α* ≈ 1, allowing the network to store the modification with high fidelity as a separate memory trace from the original version (Fig. 5D).

**Figure 6:**
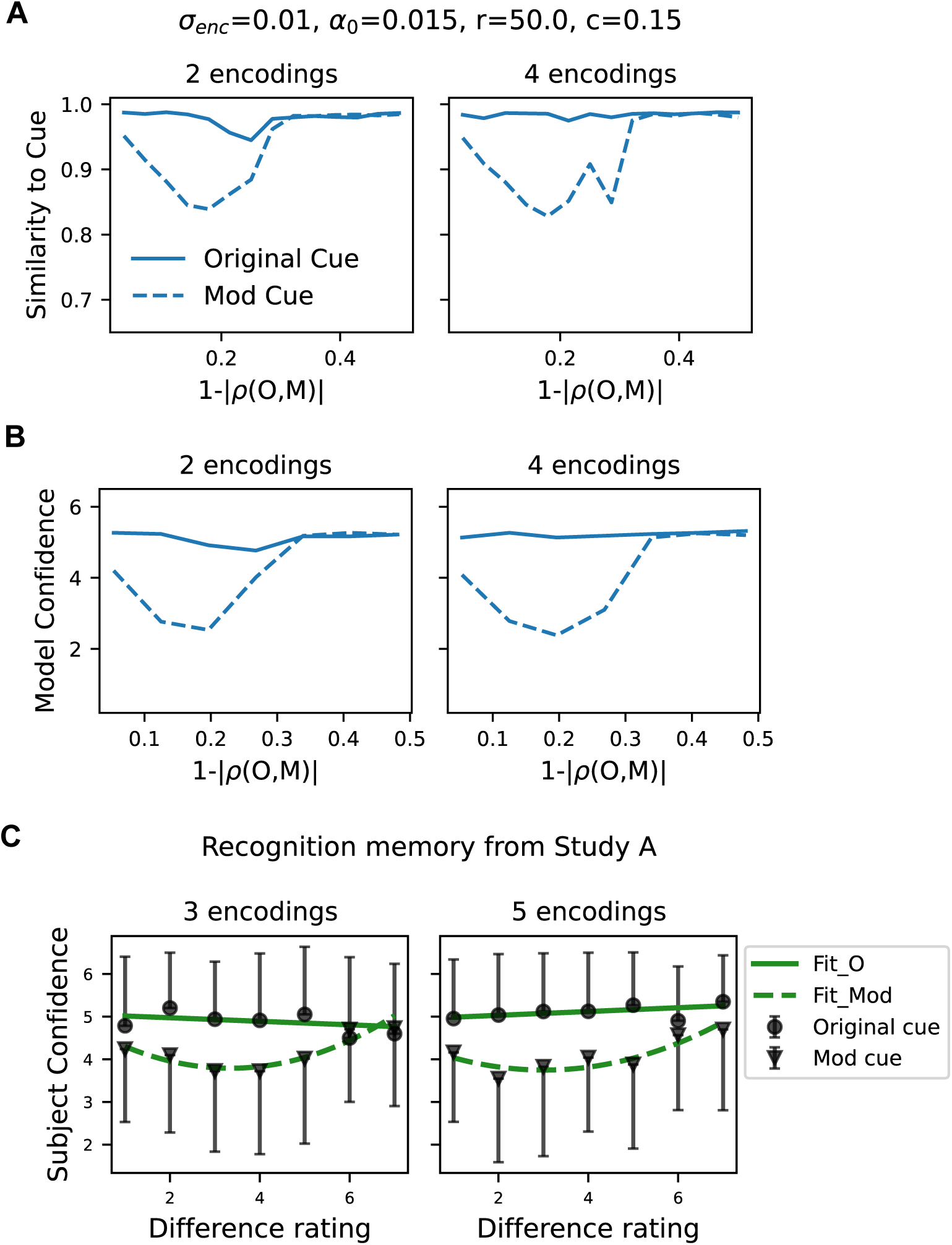
Reduced encoding strength of repeated patterns reproduces behavioral effects of encoding frequency on the recognition of original and modified stimuli. Model retrieval similarity **A** and recognition confidence **B** show robust retrieval of the original patterns (solid lines) with a small advantage after 4 (right) vs. 2 (left) encodings, while recognition accuracy of the modified pattern (dashed lines) exhibits a u-shaped pattern, replicating the behavioral findings shown in the next panel. **C**: Results of the recognition memory test in Study A after 3 (left) or 5 (right) presentations of the original stimulus.

Our results demonstrate that the competition between updating and differentiation emerges naturally in the dynamics of a Hopfield network and account for the U-shape of the recognition curve. In addition, the encoding strength modulation with PE is necessary to account for the higher robustness of memories of original versions to PE and overall weaker memory for modified versions. Our modeling results, therefore, suggest that modulation of encoding strength might be a feature of the neural mechanism underlying episodic memory in human participants.

### Accounting for the Effects of Repeated Encoding

Study A also examined the effects of repeated encodings of the original stimuli. There was a significant difference in the memory of unchanging dialogues when presented once and three times (accuracy of 1 encoding: *M* = 0.64*, SD* = 0.21; 3 encodings: *M* = 0.86*, SD* = 0.15; *p <* 0.001), but the benefit of two additional encodings diminished between three and five encodings (5 encodings: *M* = 0.90*, SD* = 0.13; *p* = 0.28). This result is in line with the logistic function we assumed for the encoding strength in the model (see Eq. (3)): If we modeled repeated storage by encoding a pattern twice in the network with the same encoding strength, the resulting attractor would become overly strong, so that future similar patterns would be prevented from being encoded in the memory. If, by contrast, the encoding strength depends on the PE, as required to reproduce Study B, then the encoding strength is quite weak (*α*_0_ ≪ 1) for a repeated pattern (Δ = 0). This is sufficient for repeated encoding to yield more precise retrieval of the original stimuli and impaired retrieval of the modified stimuli, since the modification increasingly converges towards the original pattern (Fig. 6). At the same time, the effect of repeated encoding is not strong enough to prevent the encoding of similar modified versions. Hence, our findings hint at low encoding strength, *α*(Δ = 0) ≪ 1, for repeated patterns.

### Cued Recall Analysis

Our key prediction is that encoding of a similar new memory interferes with previous memory (updating), whereas dissimilar ones are stored separately (differentiation). This prediction can be further tested experimentally by examining the contents of the cued recall test (see Methods). If the modification was very similar to the original, i.e., low difference rating, participants rarely (*<* 20%) recalled two distinct versions in studies A (Fig. 7A) and B (Fig. 7B). However, as the difference between the two versions increased, the probability of recalling two versions steadily rose to *>* 60%. This increase was confirmed by a logistic regression analysis, which showed a significant effect of difference rating on the likelihood of recalling two versions (Study A: *β* = 1.86, *p <* 0.001; Study B: *β* = 2.8709*, p <* 0.001). These experimental results confirm that when two versions of an event differ only slightly, participants tend to retain a single memory, whereas for versions that differ to a large extent, they retain two memories — as predicted by our model.

**Figure 7:**
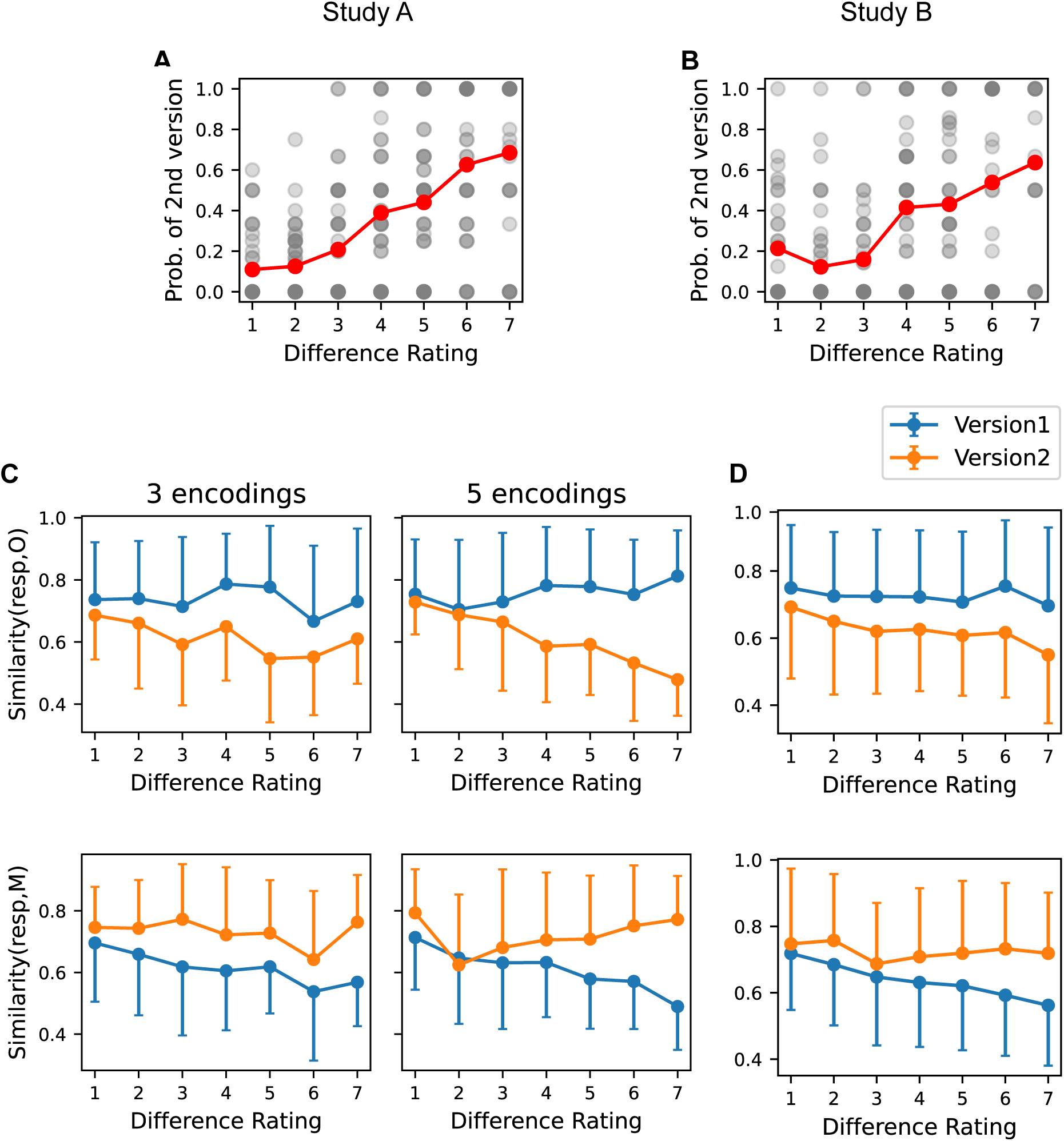
Cued recall studies show more differentiation of memory and greater separation between the remembered versions of diverging events. **A, B**: Probability of recalling two version as function of the participant-rated difference in study A and B, respectively. As the difference between original and modified conversations increases, participants are more likely to recall two versions. **C, D**: Semantic similarity between the recalled version(s) and the original (top) and modified (bottom) conversation for study A and B, respectively. Blue lines indicate the similarity of the first reported version, orange lines show the similarity of the second reported version (if given). Note that the first response is more similar to the original, and the second aligns more with the modified version.

To analyze the responses in the cued recall task, we adopted a large language model (see Methods). First, we validated the language model on our conversation data by computing the semantic similarity between original and modified versions and comparing it to the participants’ difference ratings. The two measures were broadly consistent (Fig. 8B). For similar conversations, the match between the two measures was good, i.e., the model similarity was high, if the difference rating was low. However, for divergent versions, the two measures did not align well (Fig. 8B). Furthermore, participants tended to give lower difference ratings more frequently than high ones (Fig. 8A). To capture individually perceived differences and to avoid artifacts by the model at high differences, we decided to evaluate the highly similar recall responses with the semantic similarity given by the language model, while continuing to use the participants’ ratings to measure the difference between original and modified versions.

**Figure 8:**
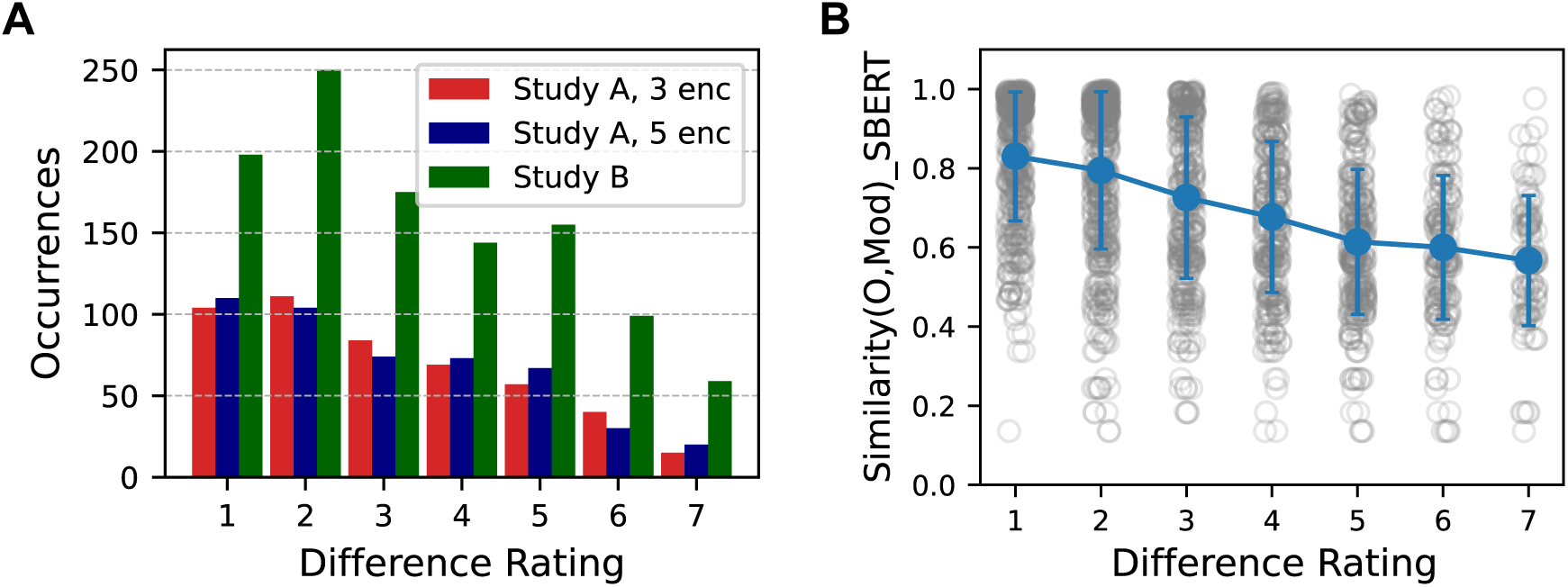
Comparison of human and large language model ratings of differences between original and modified conversations. **A**: Participants’ difference ratings for study A (3 and 5 encodings of the original) and for study B. **B**: Similarity between original and modification measured by the text embedding model approximately tracks participants’ difference ratings, except for low model similarities, which do not correlate well with participants’ ratings.

To investigate how memory responses aligned with either the original or modified dialogues, we analyzed the semantic similarity between recalled responses on the one hand, and the original or modified dialogues on the other hand. We conducted linear mixed-effects model analyses with semantic similarity as the dependent variable, and difference ratings and encoding frequency as fixed effects, including a random intercept for participants. We found that the first response was generally more similar to the original dialogue (Study A:*β* = −0.081*, p <* 0.001; Study B:*β* = −0.036*, p* = 0.009), while the second response more closely resembled the modified version (Study A:*β* = −0.1*, p <* 0.022, Study B:*β* = −0.0566*, p <* 0.139; Fig. 7C, D). These results demonstrate that participants tended to recall the original version first, followed by the modified version, suggesting that the initially encoded memory forms a stronger attractor state. This finding lends further support to the prediction of our model that encoding strength is higher for the original than the modified pattern. Moreover, we observe an increasing difference between the two recalled versions as the difference rating between the encoded conversations increased. While the first reported version remained highly similar to the original across difference levels, the second version became progressively less similar to the original as the difference rating increased. This finding supports our hypothesis that memory attractors diverge as differences between experiences become more pronounced.

## Discussion

This study examined the dynamics of competing memory representations and the role of prediction errors (PEs) in shaping their interaction. We introduced a Hopfield network model in which stimuli are encoded sequentially, with encoding strength modulated by their distinctness from previously stored patterns, which corresponds to the PE. This approach allowed us to replicate key behavioral findings from recognition and recall memory tasks, demonstrating the value of PE-modulated network dynamics in governing memory updating and differentiation.

In contrast to the classical Hopfield network model, our model stores the patterns sequentially and modulates the encoding strength of the current pattern based on the PE. The mechanism we proposed in our model gives rise to the following implications. First, a presented stimulus evokes the retrieval of a related stored memory, which is then used to modulate the encoding of the current stimulus. This mechanism reflects the findings that related memories are reinstated in the presence of current experiences (McKenzie & Eichenbaum, 2011; Nadel et al., 2012; Schlichting & Preston, 2015) and shape the encoding of new content (Gershman et al., 2013; Zeithamova et al., 2012). Second, in line with predictive coding, we showed that for our model to account for the experimental data, the strength with which a new pattern is encoded in the network must be inversely related to how predictable it was. That is, the more predictable a pattern is (lower PE), such as when a pattern is repeated, the weaker the encoding strength. Such a variation in encoding strength is consistent with observations that synaptic plasticity is higher when a novel stimulus is encoded, and neural activity is significantly reduced when a familiar stimulus is encountered (Auksztulewicz & Friston, 2016; Grill-Spector et al., 2006; Grotheer & Kovács, 2016; Summerfield et al., 2008). This agrees with our experimental observation that there is a high difference in performance when a stimulus is presented once vs. three times, with a much lower difference between three and five presentations. Third, the increasing encoding strength with increasing PE relates to observations that higher PEs can enhance synaptic plasticity, potentially via increased attentional allocation (Friston, 2010; Itti & Baldi, 2009; Ransom et al., 2017; Torrents-Rodas et al., 2021, 2023).

In contrast to findings that the PE is not only modulated by the divergence between expected and actual outcomes, but also by the strength (Chen et al., 2015; Long et al., 2016) or precision (Den Ouden et al., 2012; Greve et al., 2017; Henson & Gagnepain, 2010; Millidge et al., 2022) of the originally encoded event, in the behavioral results, we do not find a significant difference in memory performance for modifications that were presented after 3 or 5 repetitions of the original. Since the encoding modulation presented in Eq. (3) was sufficient to qualitatively replicate the behavioral data, we did not include any further modulation. In the current model, when two memories interfere, strengthening one necessarily weakens the other, which stands in conflict to other observations (Henson & Gagnepain, 2010; Long et al., 2016). However, a modulation of the encoding by the precision or strength of the original memory is possible. In our model, encoding strength follows a logistic function of PE magnitude. Although we do not explicitly model prediction precision, it could be incorporated by adjusting the slope (*r*) or midpoint (*c*) of the logistic function. A lower *r* results in a shallower, more linear encoding curve, potentially reflecting low precision or uncertain predictions, while a higher *r* leads to a sharper transition, possibly reflecting precise or confident predictions. Similarly, varying the midpoint *c* shifts the PE magnitude at which the encoding of a new memory becomes substantial: lower values of *c* imply high sensitivity even to small violations, whereas higher values require larger PEs before a modification is strongly encoded. Future work could investigate how the modulation of encoding by precision, certainty, or memory strength influences memory competition.

While our model demonstrates the role of network dynamics in explaining the competition between memories, it does not compute PEs internally; we treat the error signal as an external modulation of encoding strength. This simplification served our primary goal — to show that attractor dynamics, when scaled by a PE, reproduce key behavioral findings. Nonetheless, there are candidates for a neural mismatch signal. For example, hippocampal area CA1 is presumed to act as a comparator between inhibitory input from CA3, where stored memories are reinstated, and excitatory external input from layer 3 of the entorhinal cortex (Bein et al., 2020; Den Ouden et al., 2012; Lisman & Grace, 2005). Additionally, the inferior frontal gyrus has been implicated in signaling episodic PEs (Jainta et al., 2024; Liedtke et al., 2025), specifically by maintaining a predictive model appropriate for a given situation (Boeltzig, Liedtke, Siestrup, et al., 2025; Fujitani et al., 2024).

A key feature of our model is the interaction of overlapping patterns, demonstrating that updating or differentiation can be achieved through network dynamics, without requiring explicit decision-making or inferential processes. By systematically varying the magnitude of differences between original and modified stimuli, our study provides a graded manipulation of prediction error size. Unlike previous studies that often focused on the presence or absence of a disruption in a binary fashion, our findings highlight how different magnitudes of the PE affect memory outcomes. When two experiences are highly similar, the model predicts the formation of a single, merged representation. This accounts for the high-confidence recognition of both original and modified versions and the low rate of recalling two versions. These results are in line with previous findings that small differences between memories often lead to updating rather than differentiation (Hebscher et al., 2023; Siestrup et al., 2022; Sinclair & Barense, 2018). The model further predicts that after small deviations, the resulting representation is strongly biased towards the original and the modified version is later mistaken for the original one, consistent with prior observations of new information being mistaken for old information (Gershman et al., 2013; Hupbach et al., 2009). As differences between experiences increase to a moderate level, the robustly encoded original memory dominates and the more weakly encoded modified version is recognized with reduced accuracy. This aligns with prior research showing that existing memory traces can temper the encoding and retrieval of new, overlapping information (Bein et al., 2020; Kuhl et al., 2010; Norman & O’Reilly, 2003), particularly when the original is strongly encoded (Winters et al., 2009). When differences between originals and modifications are large, the model predicts the formation of distinct representations for each version. The increased encoding strength associated with larger deviations supports this differentiation, resulting in high recognition accuracy and an increased recall rate for both versions. This is consistent with evidence that large PEs can drive the formation of new memory representations (Bein et al., 2023; Brunec et al., 2020). Our findings are also well-aligned with the empirical finding that larger PEs promote memory for both the original and the new episode as well as source memory for the new episode, while keeping neural representations of the original information more stable (Boeltzig, Liedtke, Siestrup, et al., 2025). The predominance of the originals over the modifications is also observed in our cued recall data, where in most cases the first reported version is more similar to the original and the second version to the modification, indicating that the original version builds a stronger attractor than the modification and is therefore recalled first. Further insights could be gained with large-scale neural recordings with high temporal resolution to capture the dynamics of neural patterns.

However, while originals are overall more robustly remembered, they are not immune to change. Behaviorally, this is reflected by the reduced recognition accuracy of originals that were followed by modifications vs. those that were not (for further details, see Boeltzig, Liedtke, and Schubotz, 2025; Liedtke et al., 2025). In addition, weakly encoded originals, e.g., when presented fewer times, are more prone to being modified by PEs. In line with this experimental result, the model also shows more deterioration of originals, when they are encoded weakly (low *α*_0_) or with lower frequency. Our model predicts that the memory trace of a singly encoded original is most susceptible to updating when PEs have intermediate magnitude (Fig. 5B), because then there is interference between the competing memories, but differentiation is not reached yet. Gershman et al., 2017 came to a similar conclusion, predicting a U-shaped relationship between memory updating and the magnitude of the prediction error. This aligns with prior work showing that PEs can destabilize existing memories, rendering them susceptible to modification when new, conflicting information is introduced (Exton-McGuinness et al., 2015; Sinclair et al., 2021). This process, often framed within the reconsolidation literature (Hupbach et al., 2007; Lee, 2009; Nader & Einarsson, 2010), suggests that transformation of an original memory occurs when it is reactivated in the presence of a mismatch between expectation and outcome, whereby the extent of the update depends on the strength and reliability of the original memory (Kim et al., 2014; Przybyslawski & Sara, 1997; Suzuki et al., 2004; Vlasceanu et al., 2018) and the persistence of the unexpected event (Gershman et al., 2017; Jainta et al., 2024; Monfils et al., 2009). It has also been reported that under certain conditions, such as when the original memory is weak or the modification is repeatedly reinforced, the modified version can even dominate recall, effectively replacing the original memory trace (Schiffer et al., 2012). Future work is needed to explore how the manipulations in this experiment could be described in our model.

In summary, our study presents a computational account of how PEs influence the interaction between competing memory representations. By implementing a Hopfield network with encoding strength modulated by PEs, we capture key behavioral signatures observed in recognition and recall tasks – specifically, the transition from memory integration to differentiation across varying degrees of stimulus similarity. These findings underscore the utility of simple network dynamics in modeling complex memory phenomena and offer a foundation for future research into the neural mechanisms underlying memory updating and competition.

## Acknowledgement

This work was funded by the Deutsche Forschungsgemeinschaft (DFG, German Research Foundation), grant 397530566, FOR 2812, P1 and P2.

